# Micro-region transcriptomics of fixed human tissue using Pick-Seq

**DOI:** 10.1101/2021.03.18.431004

**Authors:** Zoltan Maliga, Ajit J. Nirmal, Nolan G. Ericson, Sarah A. Boswell, Lance U’Ren, Rebecca Podyminogin, Jennifer Chow, Yu-An Chen, Alyce A. Chen, David M. Weinstock, Christine G. Lian, George F. Murphy, Eric P. Kaldjian, Sandro Santagata, Peter K. Sorger

## Abstract

Spatial transcriptomics and multiplexed imaging are complementary methods for studying tissue biology and disease. Recently developed spatial transcriptomic methods use fresh-frozen specimens but most diagnostic specimens, clinical trials, and tissue archives rely on formaldehyde-fixed tissue. Here we describe the Pick-Seq method for deep spatial transcriptional profiling of fixed tissue. Pick-Seq is a form of micro-region sequencing in which small regions of tissue, containing 5-20 cells, are mechanically isolated on a microscope and then sequenced. We demonstrate the use of Pick-Seq with several different fixed and frozen human specimens. Application of Pick-Seq to a human melanoma with complex histology reveals significant differences in transcriptional programs associated with tumor invasion, proliferation, and immuno-editing. Parallel imaging confirms changes in immuno-phenotypes and cancer cell states. This work demonstrates the ability of Pick-Seq to generate deep spatial transcriptomic data from fixed and archival tissue with multiplexed imaging in parallel.

## INTRODUCTION

Although tissues have long been imaged using chemical stains, immunohistochemistry (IHC), and *in situ* hybridization *(1*), the introduction of single-cell RNA sequencing (scRNA-seq) has revealed an unexpected diversity of cell types and states *(2, 3*). Recently announced tissue atlases *(4–6*) combine multiplexed tissue imaging *(7–9*) with transcriptomics *(10*) to enable joint molecular and morphological analysis of human and animal tissues. Because the molecular programs that specify tissue architecture are of inherent interest, and tissues are mixtures of many cell types, there is long-standing interest in subjecting specific regions of a tissue to RNA and DNA sequencing. The earliest approaches involved manual tissue dissection using needles *(11*) but Laser Capture Microdissection (LCM)*(12*) was the first widely used approach. In LCM, an infrared laser melts an ethylene-vinyl acetate layer onto selected regions of tissue, allowing cells in that region to be recovered and processed for sequencing. More recent spatial transcriptomics approaches *(10, 13*) have made it possible to interrogate gene expression at near single-cell resolution.

Most methods for spatial transcript profiling require or work far better with, fresh frozen samples *(13*) for the simple reason that fixation damages nucleic acids. Frozen sections (those mounted in optimal cutting temperature medium; OCT) are used for intra-operative patient management, but in both pre-clinical and clinical settings formaldehyde-fixed paraffin-embedded (FFPE) specimens are more common for multiple reasons. FFPE sections are the diagnostic standard in clinical pathology, with pathology services in many teaching hospitals processing >10^5^ specimens/year; morphology is also better preserved in FFPE than OCT specimens. Archives of FFPE tissue exist for many diseases and multicenter clinical trials prefer FFPE specimens because fixed samples are easily stored and exchanged. Finally, many specimens cannot be allocated for use in research studies until they have undergone microscopic evaluation for diagnostic purposes, for example, to exclude the possibility of invasive disease. Thus, a substantial need exists for transcript profiling methods that are optimized for use with FFPE tissue in conjunction with multiplexed imaging of the same specimen. Remarkably, sequencing of single cells has recently been demonstrated using micro-regions of FFPE subjected to LCM*(14*). This result inspired us to develop approaches to micro-dissection and sequencing that were compatible with multiplexed immunofluorescence-based tissue imaging.

In this paper, we describe a method for isolating and sequencing micro-regions of interest (mROIs) from tissue guided by imaging of the same or serial (adjacent) tissue sections. The Pick-Seq approach evolved from simple and robust methods for mechanical recovery of tissue for sequencing *(15*) with the integration of compact robotic manipulators and automated mechanical recovery of tissue samples. Whereas many multiplexed imaging technologies focus on small fields of view *(8, 9*), our approach *(7*) involves whole-slide, multiplexed, subcellular resolution imaging of specimens up to several square centimeters in area; this generates single-cell imaging on a scale sufficient to provide physiological context for mROIs *(16*). Whole slide imaging makes it possible to analyze tissue structures of a wide range of spatial scales and is regarded by the FDA as a diagnostic necessity *(17*).

## RESULTS

The Pick-Seq method uses immunofluorescence whole-slide imaging of a standard 5-10 μm thick section followed by aspiration of an mROI into a liquid-filled 40 μm bore needle using a robotic arm (“picker”) and subsequent deposition into a PCR tube containing lysis buffer (**Fig. 1**) *(18*). Successful deposition can optionally be confirmed by imaging through a flat-bottom tube. To capture and sequence RNA from the mROIs, lysed cells were de-crosslinked, mRNA purified using Oligo(dT) beads, and sequencing libraries then prepared (Methods). Pick-Seq has been implemented in a commercial instrument *(19*) (RareCyte CyteFinder®) that integrates a high-resolution, slide-scanning fluorescence microscope with a robotic picker but could be performed with other microscopes and robotic manipulators.

**Fig. 1:**
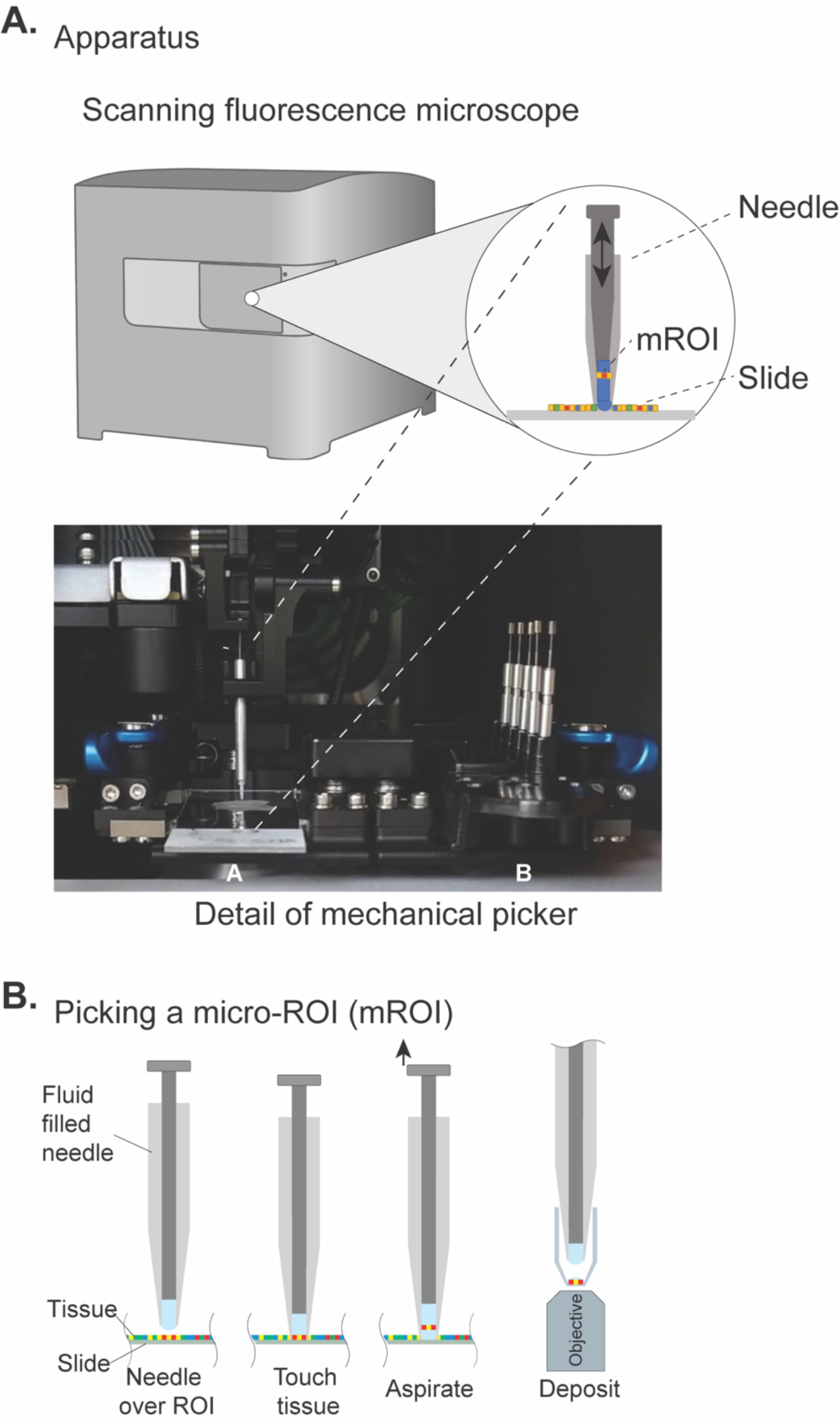
Pick-Seq technology and retrieval of micro-regions from tissue sections. **(A)** External drawing of the RareCyte CyteFinder fluorescence slide scanning microscope (left), a schematic of the picker needle (middle), and a photograph of a needle mounted on a robotic arm above a target slide in stage position A and spare needles and target tubes in position B (right). **(B)** Schematic of mROI retrieval. A fluid-filled needle attached to a robotic arm recovers tissue from a mROI. Sample recovery can be confirmed by imaging the PCR tube if desired.

To test Pick-Seq on a well-characterized tissue, we stained a section of FFPE human tonsil with B- (CD20) and T- (CD3, CD4, CD8) cell markers and, from an adjacent section, collected five mROIs from two B-cell follicles and five mROIs from an inter-follicular region rich in T cells (“T-cell zone”) (**Fig. 2, A and B**); subsequent post-pick imaging of the picked section showed that each picked mROI contained ∼5-10 cells. The resulting bar-coded RNA-sequencing (RNA-seq) libraries detected, on average, 2.700 genes per mROI, and 13,033 unique genes across all ten mROIs (**Table S1**, **Fig. 2C**). Differentially expressed genes (DEGs) between the T-cell zone and B-cell follicles included the T- and B-cell lineage markers *CD3D* and *CD19*, respectively, as expected (FDR < 0.05; **Fig. 2D**). We used CIBERSORT to deconvolve the cell composition of the mROIs from their gene expression profiles *(20*) and identified a high abundance of T or B cells expected within each compartment (**Fig. 2E**), but transcripts from other cell types were also detected (note that relative gene counts and CIBERSORT fractions are not identical because deconvolution uses gene set signatures, not single genes). Fluorescence microscopy confirmed the presence of some B cells in T-cell zones and T cells in B-cell follicles (**Fig. 2F**), but principal component analysis (PCA) was able to resolve the two B-cell follicles from each other as well as from the T-cell zone (**Fig. 2G**). When considering only the B-cell follicles, DEGs included *JCHAIN*, *MZB1*, and *CD21* (**Fig. 2H**), and subsequent imaging confirmed spatially restricted expression of CD21 (**Fig. 2B**), consistent with the absence of a follicular dendritic cell network in the sectioned plane of one follicle *(21*). We conclude that Pick-Seq can uncover spatially restricted gene expression patterns from FFPE tissue and these patterns can be confirmed by imaging.

**Fig. 2:**
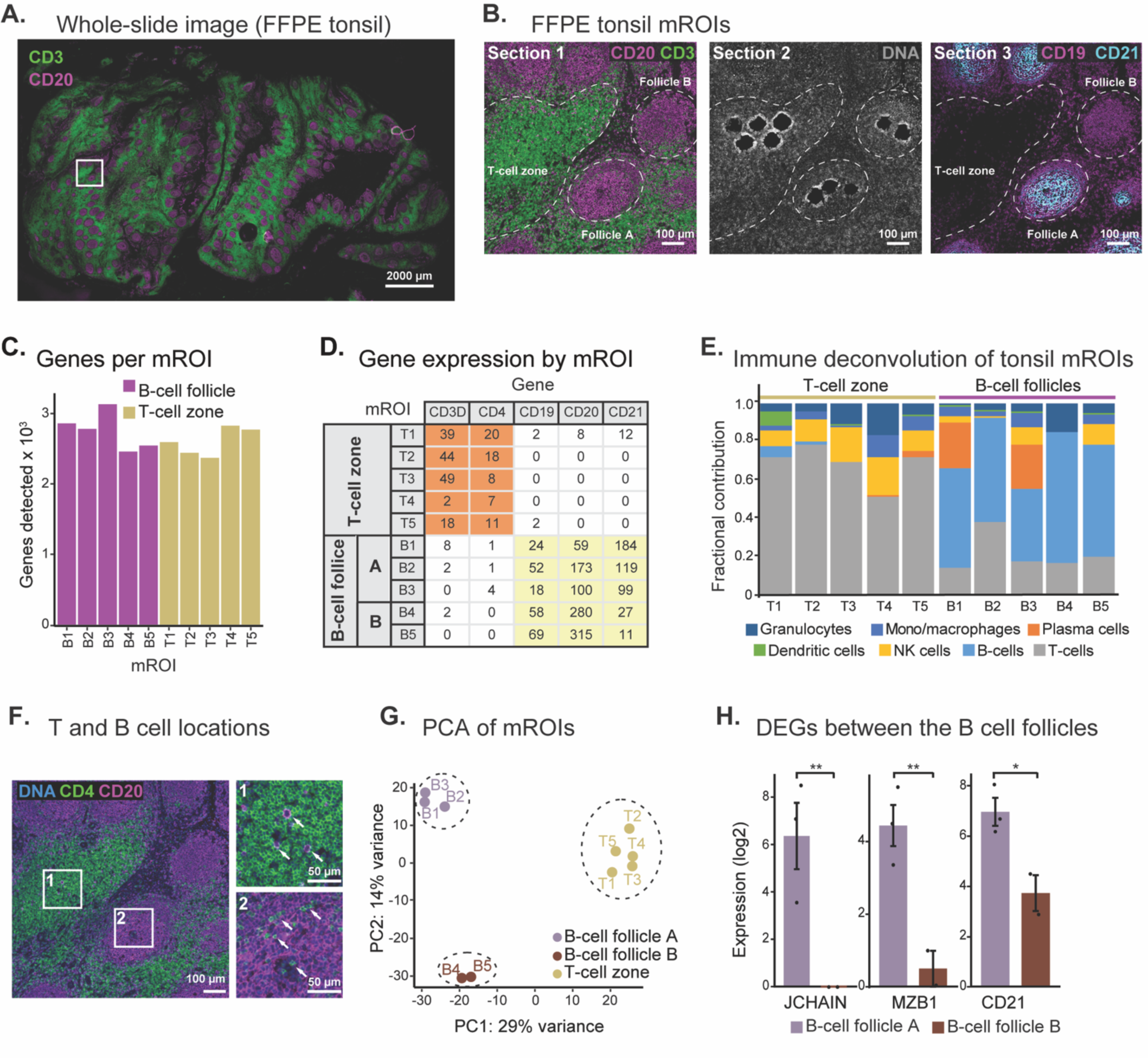
Pick-Seq analysis of an FFPE tonsil tissue section. **(A)** Whole slide fluorescence microscopy of a 5 µm thick FFPE tonsil section stained for T (CD3: green) and B (CD20: violet) cells. Scale bar, 2000 µm. **(B)** Inset region from Fig. 2A (left); corresponding region of the adjacent 5 µm thick FFPE tissue section after mROI recovery, stained to visualize nuclear DNA (DR: gray) (center); the next adjacent section was stained for a lineage marker of B cells (CD19: violet) and tonsil follicular DEG (CD21: cyan) (right). Histologic features in tonsil that guide picking are outlined (dotted lines). Scale bars, 100 µm. **(C)** Number of genes detected in each mROI extracted from FFPE tonsil. **(D)** Expression (transcripts per million; TPM) of selected cell lineage genes in each mROI. **(E)** Immune cell deconvolution for each tonsil mROI transcriptome using CIBERSORT LM22 signature. **(F)** Left, fluorescence microscopy of FFPE tonsil stained for DNA (blue), helper T cells (CD4: green), and B cells (CD20: purple). Scale bar, 100 µm. Right, insets 1 and 2. Arrows indicate scattered B-cells in T-cell zone (top) and T cells in B-cell follicle (bottom). Scale bars, 50 µm. **(G)** Principal component analysis (PCA) of tonsil mROI transcriptome data colored by histologic feature: B-cell follicle A (violet), B-cell follicle B (brown), and T-cell zone (yellow). **(H)** Expression of selected DEGs (JCHAIN, MZB1, and CD21) in mROIs from B-cell follicles A (n=3; violet) and B (n=2; brown). Data is mean ± SEM. *P<0.01; **P<0.05.

Frozen tissues have less RNA degradation than FFPE specimens, and, if available, allow for deeper RNA sequencing than FFPE. We performed Pick-Seq on a frozen section from an ER/PR double-positive breast cancer biopsy that had been stained to identify tumor (cytokeratin) and T (CD3, CD8) cells. Fourteen mROIs were recovered from three tissue microenvironments: cancer cells alone (4 mROIs), T cells outside the tumor (5 mROIs), and regions containing tumor-infiltrating T lymphocytes (TILs; 5 mROIs). In the case of TIL mROIs, imaging showed that each pick contained one (mROIs TIL1-3) or three (mROI TIL4) T cells along with an estimated four to six cancer cells (**Fig. 3A**). The RNA-seq libraries prepared from these picks detected, on average, 4,640 unique genes per mROI (∼16,800 genes in all mROIs; **Table S2**, **Fig. 3B**). We used single-sample gene set enrichment analysis (ssGSEA) to identify classes of genes that were over-represented, and we confirmed enrichment of breast cancer-related and T-cell signatures in tumor and T-cell containing mROIs, respectively (**Fig. 3C**). DEGs included estrogen receptor 1 (*ESR1)* and progesterone receptor (*PGR*; **Fig. 3D**), consistent with an ER^+^/PR^+^ luminal A tumor subtype). *ANXA1*, a gene that promotes Th1 differentiation of T cells, was found only in regions of tumor-infiltrating T cells (TILs; **Fig. 3D**), suggesting tumor specific immune-suppression *(22*). Based on these data, we conclude that Pick-Seq can identify a single T cell in a background of transcriptionally unrelated cells, although signal improves when an mROI contains three rather than one T cell (compare TIL4 to TIL1, for example; **Fig. 3C**).

**Fig. 3:**
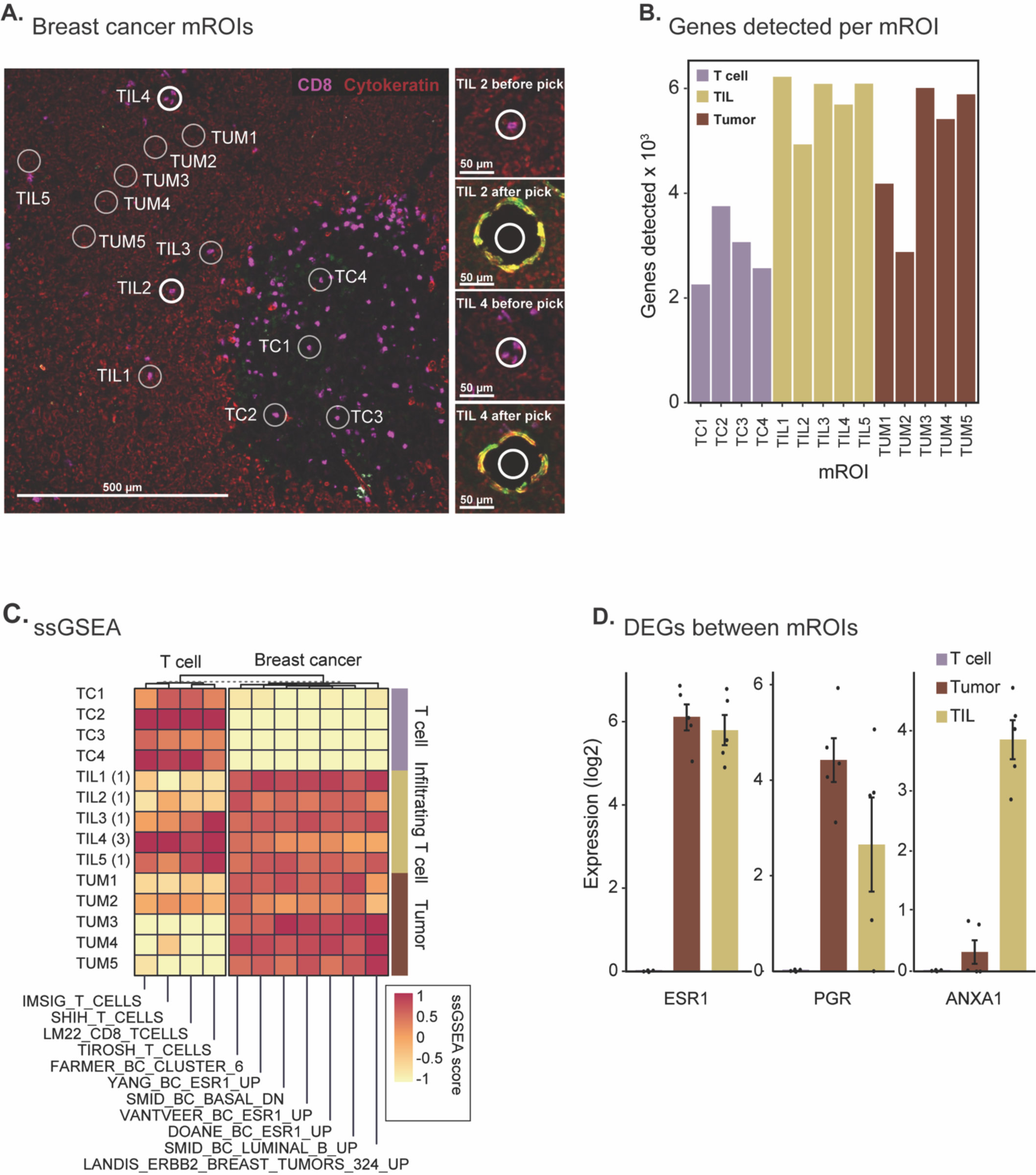
Pick-Seq of a frozen breast cancer surgical biopsy specimen. **(A)** Fluorescence microscopy of a 10 µm thick frozen section of breast cancer surgical biopsy (left) stained for tumor (cytokeratin: red) and cytotoxic T cells (CD8: magenta). Numbered mROIs are indicated as tumor (TUM), T cell (TC), and TIL. Scale bar, 500 µm. Magnified mROI containing regions before and after sample recovery of TIL2 and TIL4 samples (right). Yellow ring is due to localized tissue compaction from picking and reflects the size of the picking needle. Scale bars, 50 µm. Number of genes detected in each mROI recovered from frozen breast cancer section. Samples are coded by tissue microenvironments: T cell (TC, purple), TIL (yellow), and tumor (TUM, brown). (**C**) Single-sample gene sample enrichment analysis (ssGSEA) of mROI-derived sequence data for breast cancer (BC) and T cell related gene signatures. Number of T cells in each TIL mROI is indicated (parentheses). ssGSEA scores highlight enrichment of breast cancer-related gene signatures in tumor mROIs and T cell related signatures in the T cell rich mROIs. **(D)** Expression of selected cancer-related genes in T-cell (n=4; purple), tumor (n=5, brown), and TIL (n=5; yellow) mROIs recovered from breast tumor sample. Data is mean ± SEM.

Comparing data from FFPE tonsil with frozen breast cancer tissue (**Fig. 2C and 3B**), we found that transcriptome coverage was 50-70% greater in frozen than fixed tissue; we interpret this as arising from the higher RNA quality of frozen samples. Because polyA tail purification was used to generate sequencing libraries, RNA degradation in mROIs from FFPE specimens resulted in read data that was strongly biased toward the 3’ ends of genes as revealed by the cumulative distribution of read positions along the length of each gene (**Fig. S1, A and B**). The fraction of aligned reads was also greater for mROIs from frozen than FFPE specimens (an average of ∼ 95% for picks from frozen breast cancer samples vs. 50-80% for tonsil and melanoma; **Fig S2A**). Despite this 3’ bias, FFPE sequencing appeared to correctly capture cellular composition, as judged by imaging performed in parallel*(23*).

To study transcriptional changes associated with differences in tumor histology that impact disease outcome, we analyzed an archival FFPE *BRAF*-wild type *NF1*-mutant melanoma. This sample contained five distinct histologic regions: (i) early-stage melanoma *in situ* (MIS) (**Fig. 4, A and B**); (ii) tumor, (iii) invasive tumor margin involving tumor growth into the normal dermis; (iv) tumor-adjacent infiltrating lymphocytes (TILs) representing a brisk TIL response (**Fig. 4B**); and (v) exophytic melanoma projecting up towards the surface of the skin (not shown). Seventy-five mROIs samples were collected from this sample, each containing approximately 5-20 cells, with an average of 2,377 genes detected per mROI (**Table S3, Fig. S2B**), consistent with results from the FFPE tonsil sample (**Fig. 2C**). The PCA landscape across all mROIs (25% of variance explained in PC1 and PC2) revealed four clusters with mROIs from exophytic and invasive tumor near each other and distinct from the MIS and brisk TIL regions (**Fig. 5A**). The mROIs from the invasive margin clustered close to, but still largely separate from, the exophytic and invasive tumor clusters. Pairwise analysis among all five histological sites identified 208 to 947 DEGs; data in **Fig. 5B** shows 705 DEGs for tumor vs. MIS regions. Tumor enriched DEGs included *S100B*, a progression marker *(24*), and *CD63*, a negative regulator of the epithelial-mesenchymal transition in melanoma *(25*) (**Fig. 5C**). Imaging of an adjacent tissue section confirmed the melanocyte-restricted expression of S100B and CD63 in tumor regions (**Fig. 5D**), consistent with the annotation of these proteins as progression markers. When Pick-Seq was repeated on adjacent specimens, batch-independent clustering by region was observed, demonstrating the reproducibility of the method (**Fig. 5, E and F**).

**Fig. 4:**
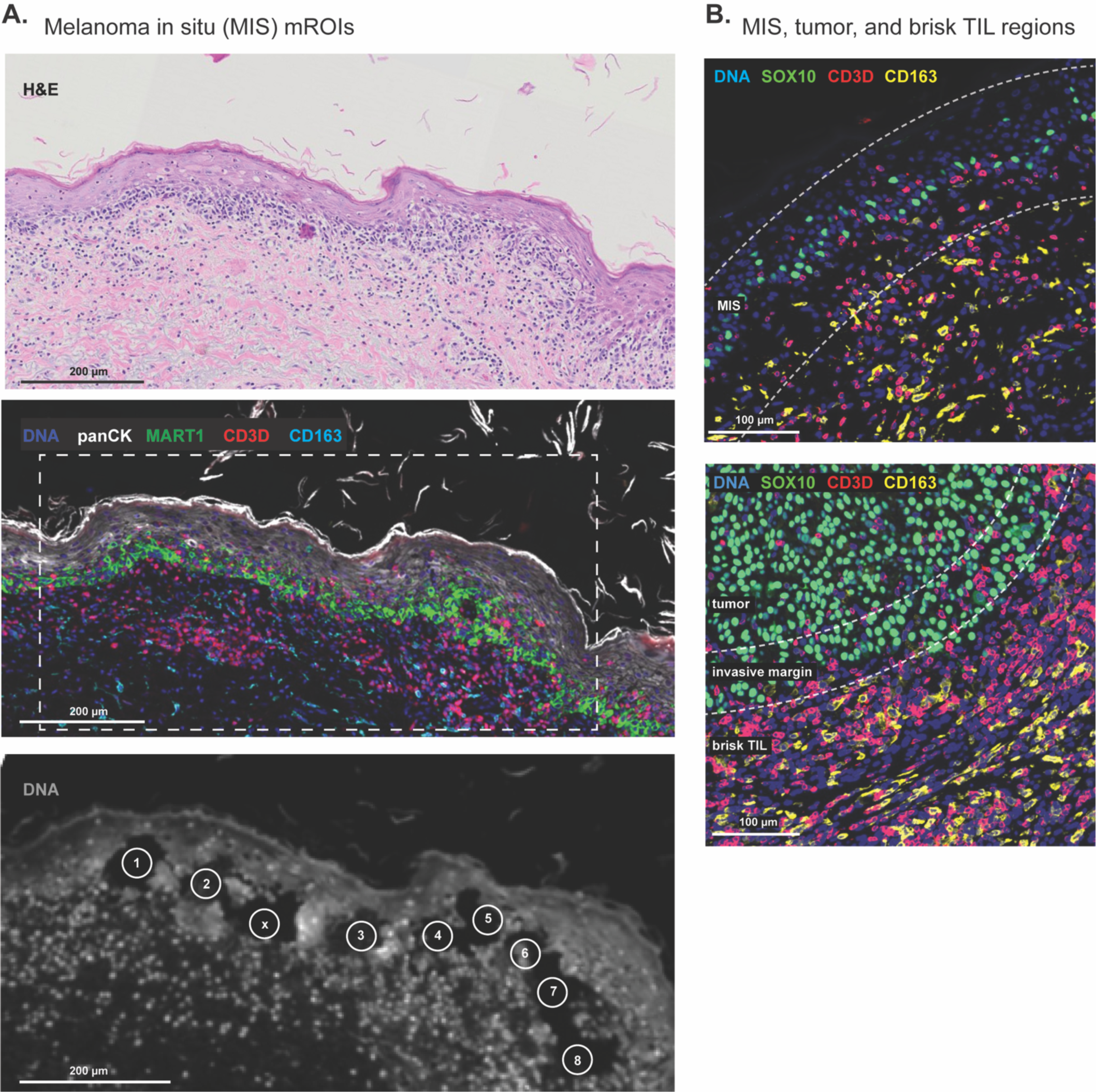
A complex archival melanoma FFPE specimen. **(A)** H&E (top panel) and fluorescence microscopy (middle panel) images of a melanoma in situ (MIS) region stained for DNA (blue), keratinocytes (cytokeratin: white), melanocytes (MART1: green), T cells (CD3D: red), and macrophages (CD163: cyan). Magnified corresponding region (bottom panel) in an adjacent tissue section stained for DNA (DR: gray). Sites of MIS mROI extraction are indicated. Scale bars, 200 µm. **(B)** Fluorescence microscopy images of histologic features in melanoma tissue stained for DNA (blue), melanocytes (SOX10: green), T cells (CD3D: red), and macrophages (CD163: yellow). Top, a region of melanoma in situ. Bottom, a region of invasive melanoma containing tumor center, invasive margin, and adjacent brisk TIL region. Scale bars, 100 µm.

**Fig. 5:**
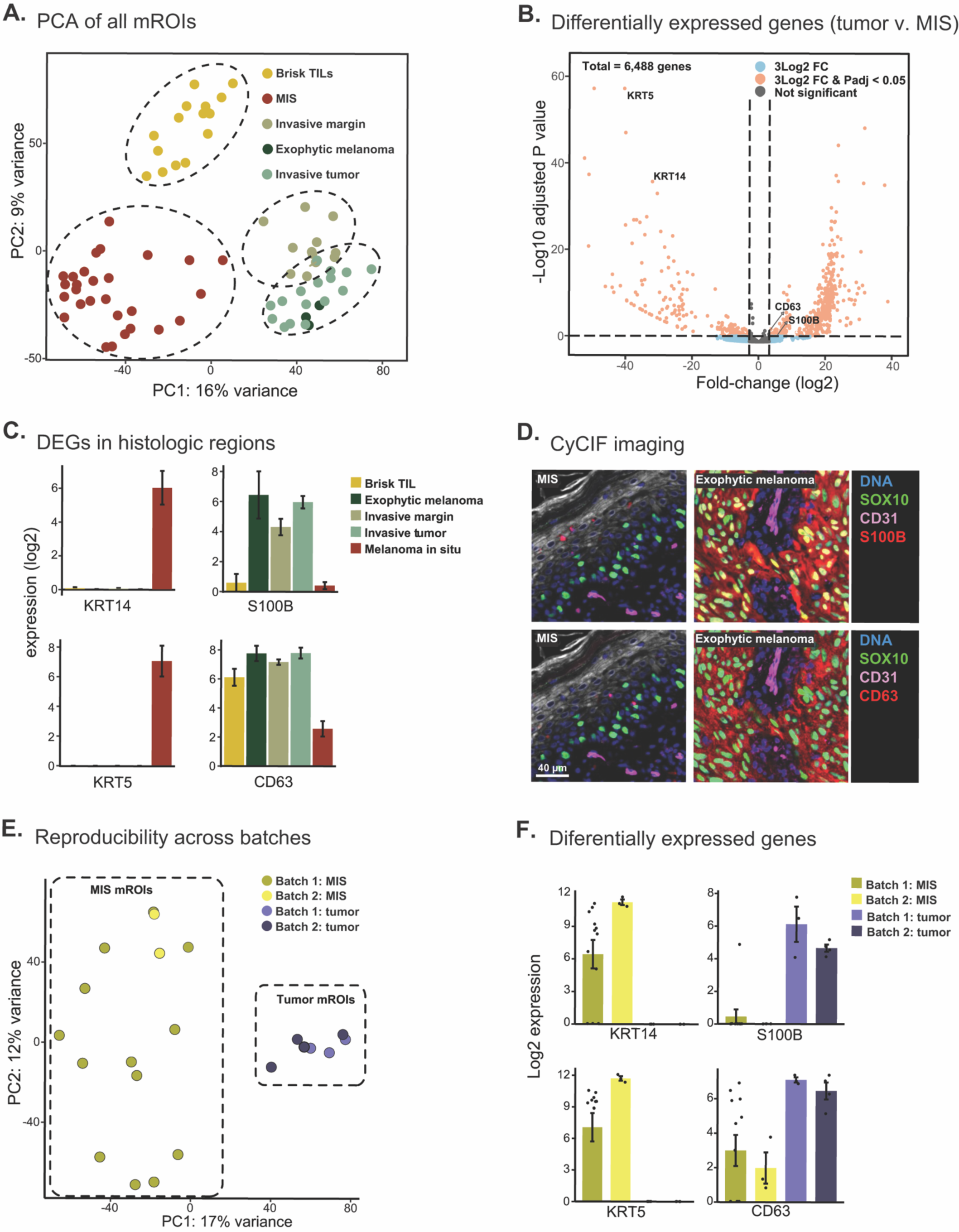
Pick-Seq of archival FFPE melanoma specimen examining transcriptional differences associated with distinct tumor domains. **(A)** Principal component analysis (PCA) of melanoma mROI transcriptomes. Colors indicate regional histopathology: brisk TIL (BTIL: yellow), MIS (red), invasive margin (IM: olive green), exophytic melanoma (EM: dark green), and center of invasive melanoma tumor (IT: light green). EM and IT are considered tumor samples in this analysis. **(B)** Fold-difference and significance for expression of 6,488 genes between tumor (n=19) and MIS (n=28) samples. DEGs above (blue) and below (orange) a significance threshold (P-adjusted = 0.05) are indicated. **(C)** Expression of selected genes (KRT14, KRT5, S100B, and CD63) in each histologic region of melanoma (same mROIs as **(A)**). Mean ± SEM for mROIs of same histology. **(D)** Fluorescence microscopy image of MIS (left) and exophytic melanoma (right) stained for DNA (blue), melanocytes (SOX10: green), blood vessels (CD31: violet), and S100B (red, top panel) or CD63 (red, bottom panel). Scale bar, 40 µm. **(E)** Principal component analysis (PCA) of mROIs retrieved from exophytic melanoma tumor (purple) or MIS regions (yellow) from different tissue sections of the same patient in separate experiments. Samples are coded by batch and histologic feature. **(F)** Expression of selected genes (KRT14, KRT5, S100B, and CD63) in exophytic melanoma tumor (purple) or MIS (yellow) mROIs. Mean ± SEM for mROIs of same histology.

Enrichment of immune-related signatures in mROIs from the TIL and MIS regions (**Fig. 6**) was consistent with imaging of an adjacent tissue section for the presence of macrophages and T cells (**Fig. 4B**). The ratio between signatures for MITF (a transcription factor) and AXL (a receptor tyrosine kinase) has been studied extensively in melanoma *(26, 27*), and in our data, GSEA demonstrated enrichment of MITF programs and downstream targets in the exophytic melanoma region as compared to MIS (**Fig. 7, A and B**); in MIS, a MITF-low, AXL-high transcriptional state was observed (**Fig. 7A**). Fluorescence microscopy confirmed higher MITF protein levels in melanocytes within the tumor region relative to MIS (**Fig. 7, C and D**), but AXL expression in MIS was restricted to the membranes of epithelial keratinocytes (**Fig. 7E**). Thus, fluorescence microscopy shows that differential AXL gene expression between MIS and tumor is a consequence of the cellular composition of the mROIs rather than a change in melanocyte biology, demonstrating the value of combining RNA expression data with protein expression data.

**Fig. 6:**
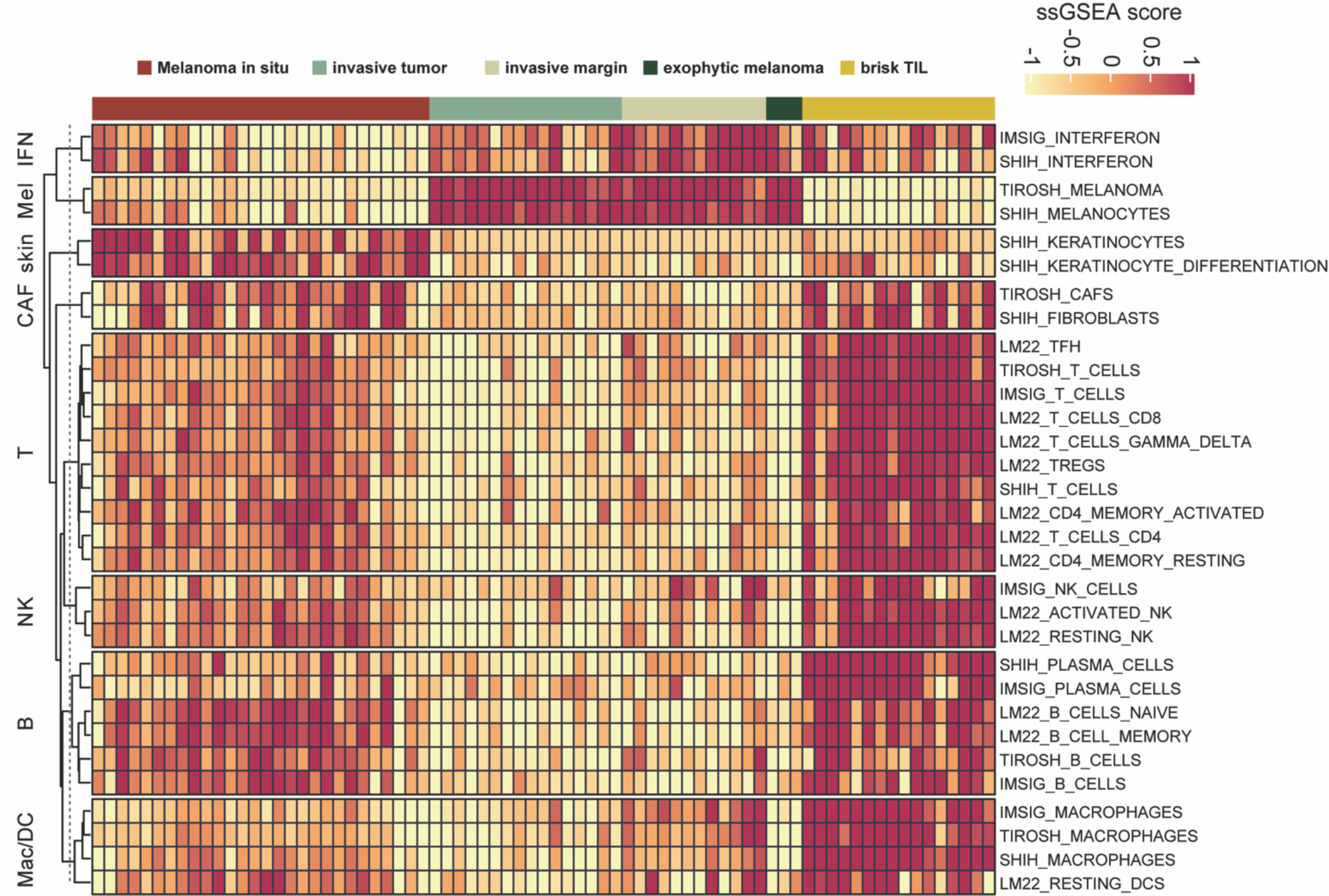
ssGSEA of immune signatures for melanoma mROIs. Single-sample gene set enrichment analysis (ssGSEA) for immune cell, skin, and melanoma-related gene signatures (rows) in mROIs of similar histology (columns) from FFPE melanoma.

**Fig. 7:**
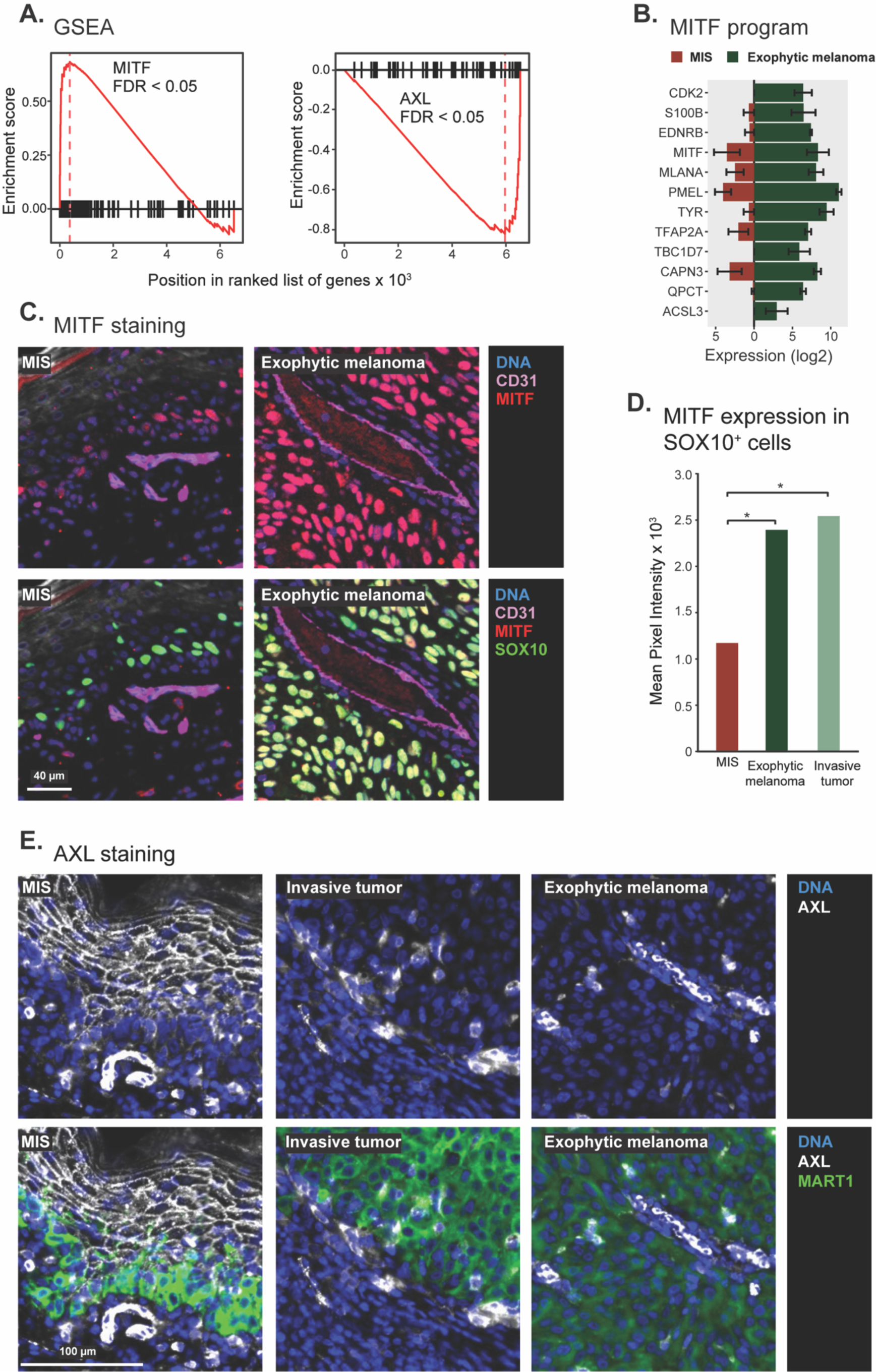
Analysis of transcriptional signatures in melanoma. **(A)** Gene set enrichment analysis for MITF pathway (left panel) and AXL pathway (right panel) in exophytic melanoma (n=3) and MIS (n=28) samples. FDR < 0.05. **(B)** Expression (log2) of selected genes in the MITF pathway in MIS (n=28; red bars to the left) and exophytic melanoma (n=3; green bars to the right) samples. Data is mean ± SEM. **(C)** Fluorescence microscopy of MIS (left) and exophytic melanoma (right) stained for DNA (blue), blood vessels (CD31: violet), and the melanoma progression marker MITF (red); the bottom panels are also stained for melanocytes (SOX10: green). Scale bar, 40 µm. **(D)** MITF protein detected by fluorescence microscopy in melanocytes (SOX10 positive cells) in MIS, exophytic melanoma, or invasive tumor regions (* P value < 0.05, t-test). **(E)** Fluorescence microscopy images of MIS (left), invasive tumor (center), or exophytic melanoma (right) stained for DNA (blue), AXL (gray), and, in the bottom panels, melanocytes (MART1: green). Scale bar, 100 µm.

NanoString GeoMx™ DSP is a leading, commercially available method for high-plex, spatial transcriptomic analysis of FFPE tissues *(28, 29*). We used it to evaluate the performance of Pick-Seq on serial sections containing regions of exophytic melanoma and MIS. GeoMx measures the abundance of ∼1,800 transcripts: 1,571 of these were detected by Pick-Seq (**Table S4**, **Fig. 8**). Most of the 94 DEGs (exophytic melanoma vs. MIS) common to GeoMx and Pick-Seq were concordantly up- or down- regulated in both assays (**Fig. 9**). Concordance between Pick-Seq and GeoMx strongly suggests that the two methods correctly capture significant differences between tissue regions. However, Pick-Seq sampled regions approximately 25-fold smaller in area than GeoMx (40 μm vs. ∼200 μm diameter) while identifying ∼2.2-fold more DEGs (FDR < 0.05; **Fig. 9** inset). The 40 μm size of Pick-Seq mROIs compares favorably to the 100 μm raster used for spatial transcriptomics on frozen specimens *(10, 30*), and the depth of sequencing and integration with imaging retains the advantages of other emerging methods involving “*high-definition spatial transcriptomics*” *(31*) while extending the application to fixed specimens.

**Fig. 8:**
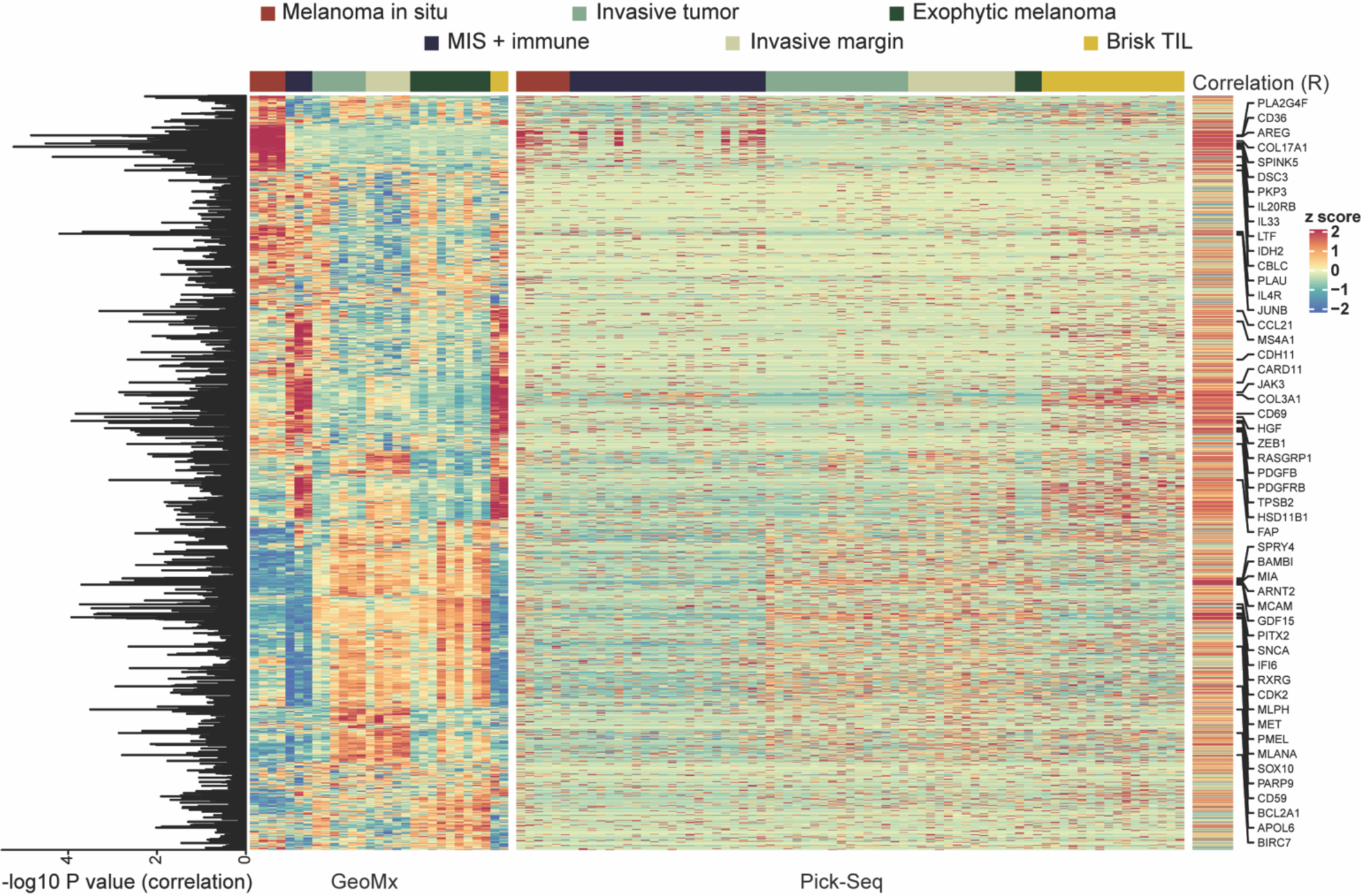
Gene expression analysis with Pick-Seq and GeoMx. Expression of 1571 genes (rows) detected by both GeoMx and Pick-Seq organized by mROI and tissue histology (columns). Correlation for each gene between the two methods (right) and significance of correlation (P-value) between the two technologies (left) are indicated.

**Fig. 9:**
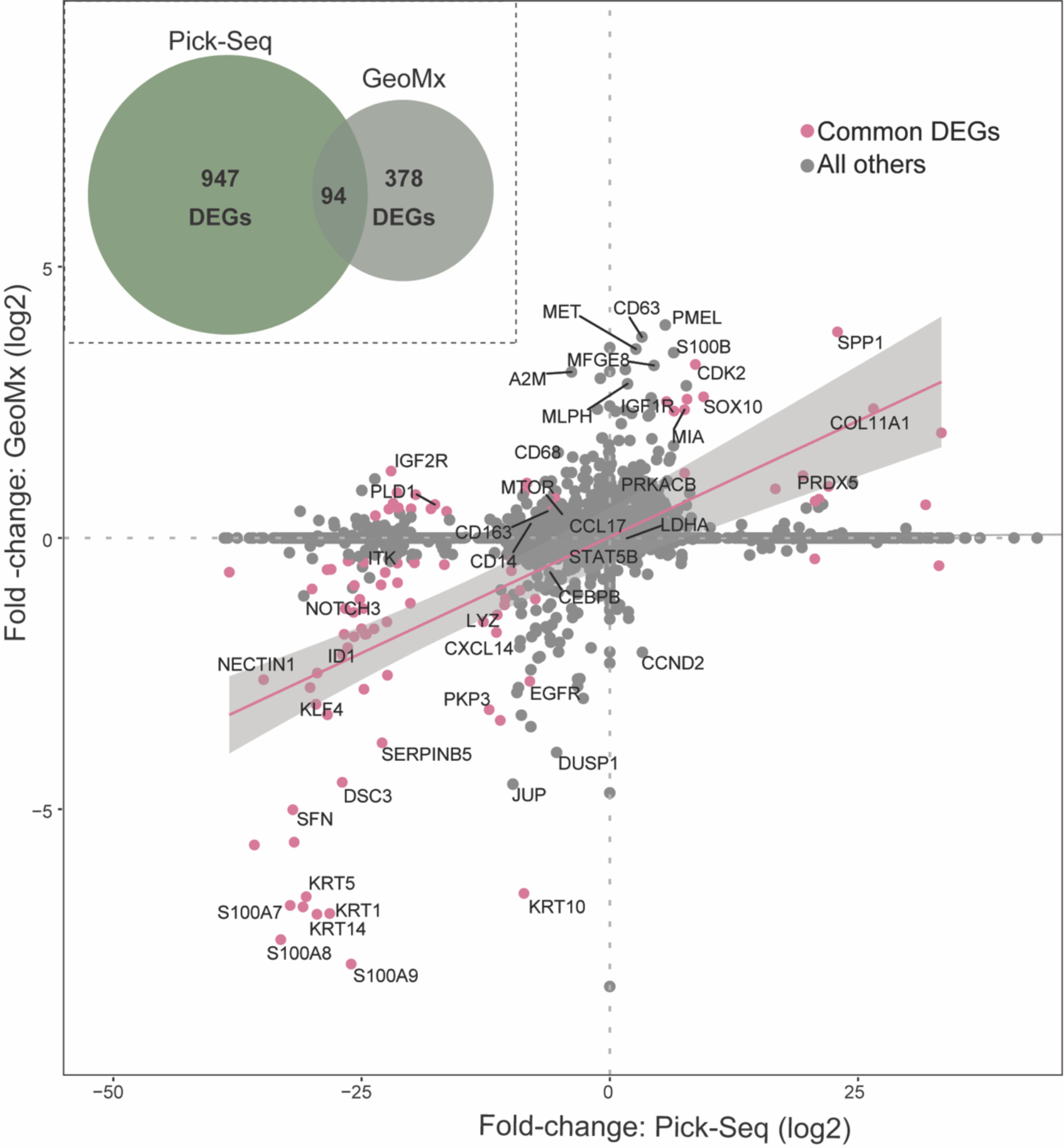
Comparison of Pick-Seq and GeoMx differentially expressed genes. Differential expression (log2) for 94 DEGs (pink) measured using Pick-Seq and GeoMx. All other genes (gray). Trend-line (pink) and confidence interval (95%, gray) are indicated. For Pick-Seq, n=3 exophytic melanoma and n=28 MIS samples. For GeoMx, n=9 exophytic melanoma and n=7 MIS samples. Inset, top left, Venn diagram indicates the number of DEGs measured using one or both technologies.

## DISCUSSION

The data in this paper establishes that micro-mechanical isolation of tissue mROIs using compact robotics integrated with multiplexed fluorescence imaging is a simple and sensitive means of spatially-resolved transcriptional profiling of fixed and frozen tissue. Pick-Seq is sufficiently reliable that in a typical use case, it proved possible to sequence 75 successive micro-regions spanning multiple tumor stages and histomorphologies in a single complex melanoma. Pairwise comparison of RNA from melanoma *in situ*, invasive tumor, and areas of active immunosurveillance identified ∼200 to 950 DEGs, which included genes known to be differentially expressed by tumor stage or extent of lymphocyte infiltration as well as newly identified genes. In our work on tumor atlases, tight integration of imaging and transcript profiling is essential, but it is also convenient for the now-common practice of validating cRNA-Seq or micro-region sequencing (mrSeq) results using immunofluorescence imaging *(32*).

Mechanical isolation avoids complex chemistry and RNA-damaging lasers, and can, in principle, be used with many different sequencing and imaging approaches; the implementation described here has been commercialized, making it readily accessible. A wide variety of methods are being developed for spatial transcriptomics of frozen specimens *(33*), but FFPE tissue presents additional challenges. These challenges merit overcoming because FFPE tissues are more widely available in research and clinical settings (including in histopathology archives), better preserve morphology and have fewer pre- analytical variables; we therefore expect Pick-Seq to have its greatest impact in the analysis of fixed and archival tissue.

In most settings, Pick-Seq is not a true “single-cell” method, but we have shown that it is possible to pick one cell of interest when it is surrounded by other cell types and deconvolve the data; with improvements in sequencing technology (e.g. use of Smart-3SEQ or Smart-seq3)*(14, 34*) and computation, this may approach single-cell resolution *(20*). In our specimens, Pick-Seq outperformed GeoMx, the leading micro-region sequencing (mrSeq) platform, with respect to spatial resolution as well as the number of DEGs, and both appear to be more sensitive than laser capture microdissection as usually deployed *(35*). Implementing different sequencing workflows, including methods for single-cell sequencing of T-cell receptors *(35*), represents a direct and potentially impactful extension of the method described here. We also anticipate continued improvement in Pick-Seq through the use of smaller needles, faster picking, and integration with whole-slide imaging to enable a straightforward comparison of molecular and morphological features of normal and diseased tissues. More generally, the integration of transcript profiling with highly-multiplexed tissue imaging on microscope platforms promises to substantially advance the goal of linking our understanding of disease genomics with histopathology for research and diagnostic purposes*(5*).

## MATERIALS AND METHODS

### Tissue procurement

Frozen breast cancer blocks were obtained from a commercial vendor (Origene, Rockville MD) and sectioned at a thickness of 10 microns by the University of Washington Pathology Core. FFPE tonsil sections were obtained from Zyagen (San Diego, CA). Freshly harvested tissue was fixed by the vendor in 10% neutral buffered formalin and processed for paraffin embedding. Paraffin blocks were sectioned at a thickness of 5 microns and mounted on positively charged slides.

FFPE melanoma sections were obtained under IRB oversight as 5 micron sections from the archives of the Dermatopathology Core at Brigham and Women’s Hospital. The sections originated from a biopsy removed from a 61 year-old male, non-smoker, with extensive sun exposure. He presented with a pigmented lesion on his forearm with recurrent intermittent bleeding that became continuous after a trauma. The patient was also diagnosed with basal cell carcinoma on the right side of the chest. The clinical pathology reported a malignant melanoma, depth of 14.0 mm, anatomic level IV, with extensive associated melanoma in situ. Superficial spreading, intraepidermal component, vertical growth phase, and ulceration were also noted as present. Non-brisk tumor-infiltrating lymphocytes were observed. There was no report of perineural or vascular invasion. The sample tested negative for BRAF V600E and PD-L1. This study was approved by the Institutional Review Board of the Harvard Faculty of Medicine (FWA00007071, Protocol IRB18-1363). A waiver of the requirement to obtain consent was deemed appropriate.

### Instrumentation

The RareCyte® imaging and picking platform performs four or six-color fluorescence imaging and has a microscope stage that employs a kinematic mount to ensure highly reproducible positioning of the slide; X–Y displacement upon reloading is approximately 2–3 μm. Scanning of each slide in four channels takes about 12 min inclusive of image plane determination. The fluid-coupled picking system positions a needle above the slide stage for retrieval of individual mROIs. The needle tip mechanically dislodges the tissue region with positive pressure into an imaging tube; visual confirmation of deposition is optional. The system is automated and does not require high technical skills; the rate of successful cell retrieval is ∼80–90%. Additional feature of the system, originally developed for isolation of circulating tumor cells, have been described previously *(19*)

### OCT tissue staining and imaging

Frozen biopsy samples were sectioned 10 µm thick in optimal cutting temperature (OCT) mounting medium, thawed (25°C, 5 minutes), dehydrated in acetone (-20°C, 10 min), rehydrated in PBS/0.25% Triton X/0.0025% RNasin Plus (Promega) (twice, 0°C, 3 min each) then washed PBS/0.0025% RNasin Plus (0°C). Prior to staining slides were blocked with ice-cold blocking buffer 1 (PBS/6% BSA/0.1% RNasin Plus) for 10 min on ice. Sections were stained with primary antibodies (**Table S5**) and SYTOX Orange (nuclear stain) diluted in a blocking buffer and incubated on ice for 30 min. After staining, slides were washed twice for 3 min in ice-cold 1X PBS/0.25% Triton X/0.0025% RNasin Plus, followed by a wash in ice-cold 1X PBS/0.0025% RNasin Plus. Stained sections were mounted with 1X PBS/0.1% RNasin Plus and scanned on a CyteFinder instrument to identify regions of interest. Coverslips were removed, and sections were dehydrated in a series of ice-cold solutions each containing 0.0025% RNasin Plus: 1X PBS for 1 min, 1X PBS for 1 min, 75% ethanol for 1 min, 95% ethanol for 1 min, 100% ethanol for 1 min. Slides were left in ice-cold 100% ethanol prior to mROI retrieval.

### FFPE tissue staining and imaging

FFPE sections were deparaffinized and rehydrated using the Histogene Refill Kit (Arcturus). Slides were immersed in xylene for 10 min, a second jar of xylene for 15 min then incubated in an ethanol series (100% ethanol for 4 min, 95% ethanol for 4 min, 75% ethanol for 4 min) followed by 1X PBS for 4 min. For antigen retrieval, slides were incubated in Leica 1X Tris/EDTA pH 9 retrieval solution at 95°C for 10 min, then washed in 1X PBS/0.25% Triton X three times (3 min each wash). To reduce autofluorescence, slides were submerged in 1X PBS with 4.5% hydrogen peroxide and 24 mM NaOH and photobleached between two light sources for 1 hr. Slides were washed twice in 1X PBS/0.25% Triton X (3 min per wash), followed by a 1X PBS wash, and stored overnight at 4°C. Melanoma sections were dewaxed and subjected to antigen retrieval using a Leica Biosystems BOND RX automated slide stainer, following the manufacture’s protocols.

Prior to staining, slides were blocked for 1 hour at room temperature with blocking buffer 2 (1X PBS/10% goat serum/6% BSA/0.1% PEG). Primary antibodies diluted in blocking buffer 2 (**Table S5**) were applied to the section and incubated for 1 hr at room temperature. After the primary antibody incubation, slides were washed two times with 1X PBS/0.25% Triton X (3 min per wash), followed by a wash with 1X PBS. Secondary antibodies diluted in blocking buffer 2, were applied to the sections and incubated for 1 hr at room temperature. Slides were then washed twice with 1X PBS/0.25% Triton X (3 min per wash) followed by a wash with 1X PBS. For the second round of staining on the same section, the above primary antibody incubation step was repeated after a blocking step. To visualize the nuclei, slides were stained with a 1:100,000 dilution of SYTOX Orange for 15 min at room temperature followed by two washes in 1X PBS/0.25% Triton X (3 min per wash). Stained sections were mounted with RareCyte mounting media and coverslip applied. Stained slides were scanned on a CyteFinder instrument.

### Pick-Seq tissue section preparation

For each FFPE tissue sample, a serial section adjacent to the immunofluorescence (IF) stained section was prepared for microregion retrieval and downstream RNA-seq analysis. Sections were deparaffinized and rehydrated using the Histogene Refill Kit. Slides were immersed in xylene for 5 min, followed by incubation in a second jar of xylene for 5 min. Slides were then incubated in a series of ice-cold solutions with 0.0025% RNasin Plus (Promega): 100% ethanol for 1 min, 95% ethanol for 1 min, 75% ethanol for 1 min, 1X PBS for 1 min, and another tube of 1X PBS for 1 min. Slides were stained with 50 µM DR,™ a Far-Red DNA Dye (ThermoFisher) in PBS, with 0.1% RNasin Plus for 2 min on ice. Sections were dehydrated in a series of ice-cold solutions with 0.0025% RNasin Plus: 1X PBS for 1 min, 1X PBS for 1 min, 75% ethanol for 1 min, 95% ethanol for 1 min, 100% ethanol for 1 min. Slides were left in ice-cold 100% ethanol prior to microregion retrieval.

### Pick-Seq micro-region retrieval

IF stained sections were evaluated to identify regions of interest for transcriptional analysis (**Table S6**). Tissue architecture of the IF-stained and DR-stained sections were compared to identify the corresponding regions of interest on the DR-stained slide. Prior to microregion retrieval, slides were removed from 100% ethanol and allowed to air dry. Slides were loaded into a CyteFinder instrument (RareCyte), and microregions were retrieved using the integrated CytePicker module with 40 µm diameter needles. Tissue microregions were deposited with 2 µl PBS into PCR tubes containing 18 µl of lysis buffer: 1:16 mix of Proteinase K solution (QIAGEN) in PKD buffer (QIAGEN), with 0.1% RNasin Plus. After deposit, tubes were immediately placed in dry ice and stored at -80°C until ready for downstream RNA-seq workflow.

### Tissue microregion lysis and mRNA enrichment

PCR tubes containing tissue microregions in the lysis buffer were removed from the freezer, allowed to thaw at room temperature for 5 min, and incubated at 56°C for 1 hr. Tubes were briefly vortexed, spun down, and placed on ice. Dynabeads Oligo(dT)25 beads (ThermoFisher) were washed three times with ice-cold 1X hybridization buffer (NorthernMax buffer (ThermoFisher) with 0.05% Tween 20 and 0.0025% RNasin Plus) and resuspended in original bead volume with ice-cold 2x hybridization buffer (NorthernMax buffer with 0.1% Tween 20 and 0.005% RNasin Plus). A volume of 20 µl of washed beads was added to each lysed sample, mixed by pipette, and incubated at 56°C for 1 min followed by room temperature incubation for 10 min. Samples were placed on a magnet and washed twice with an ice-cold 1X hybridization buffer, then once with ice-cold 1X PBS with 0.0025% RNasin Plus. The supernatant was removed, and the pellet was resuspended in 10.5 µl nuclease-free water. Samples were incubated at 80°C for 2 min and immediately placed on a magnet. The supernatant was transferred to new PCR tubes or plates, and placed on ice for subsequent whole transcriptome amplification or stored at -80°C.

### Whole transcriptome amplification, RNA-seq library preparation, and sequencing

Reverse transcription and cDNA amplification were performed using the SMART-Seq v4 Ultra Low Input RNA Kit for Sequencing (Takara Bio, Kusatsu, Shiga, Japan). The resulting amplified cDNA libraries were assessed for DNA concentration using the Qubit dsDNA HS Assay Kit (ThermoFisher) and for fragment size distribution using the BioAnalyzer 2100 High Sensitivity DNA Kit (Agilent). The fragment size distribution is used to determine how to make the cDNA into a sequencing library. Tissues, where the majority of cDNA was >500 bp in length, was prepared into a library using Nextera tagmentation method (Illumina) while if the majority of the cDNA was <500 bp in length the cDNA was cleaned up and adapters were ligated following the ThruPLEX method (Takara Bio).

cDNA from tonsil and breast cancer tissue microregion samples were prepared into sequencing libraries using Nextera XT library preparation (Illumina), while cDNA from melanoma microregion libraries were prepared with ThruPLEX DNA-seq Kit (Takara Bio). The resulting libraries were quantitated using the Qubit dsDNA HS Assay Kit and quality was assessed on a BioAnalyzer 2100 High Sensitivity DNA Kit. Libraries were pooled at equimolar ratios and sequenced in-house using an Illumina MiSeq or on an IlluminaNextSeq at Biopolymers Facility at Harvard Medical School.

### RNA-seq data processing

The raw FASTQ files were examined for quality issues using FastQC (http://www.bioinformatics.babraham.ac.uk/projects/fastqc/) to ensure library generation and sequencing are suitable for further analysis. The reads were processed using the bcbio pipeline v.1.2.1 software *(36*). Briefly, reads were mapped to the GRCh38 human reference genome using HISAT2 *(37*) and Salmon *(38*). Length scaled transcripts per million (TPM) derived from Salmon was passed through the ARSeq pipeline v.2.2.14 (https://github.com/ajitjohnson/arseq) for downstream analysis. The DESeq2 R package *(39*) was used to generate the normalized read count table based on their estimateSizeFactors() function with default parameters by calculating a pseudo-reference sample of the geometric means for each gene across all samples and then using the “median ratio” of each sample to the pseudo-reference as the sizeFactor for that sample. The sizeFactor is then applied to each gene’s raw count to get the normalized count for that gene. DESeq2 was also used for differential gene expression analysis. A corrected P value cut-off of 0.05 was used to assess significant genes that were up-regulated or down- regulated using Benjamini-Hochberg (BH) method. The combat function from the R package SVA was used to account for batch effects when combining two independent Pick-Seq experiments for assessing technical reproducibility.

### Pathway enrichment analyses

A compendium of biological and immunological signatures was identified from publicly available databases or published manuscripts for performing enrichment analysis. To perform gene set enrichment analysis, two previously published methods (Gene Set Enrichment Analysis (GSEA) *(40*) and single- sample GSEA (ssGSEA)) were primarily used. The R package clusterProfiler *(41*) was used to perform GSEA and the R package GSVA *(42*) was used to perform ssGSEA which calculates the degree to which the genes in a particular gene set are coordinately up- or down-regulated within a sample. The breast cancer signatures were curated from MsigDB *(43*), and immune cell-related and melanoma- related (MITF pathway and AXL pathway) signatures were curated from published studies *(32, 44–46*).

### CIBERSORT analysis

The normalized counts’ table was inverse log2 transformed and uploaded to the CIBERSORT web app v1.06 (https://cibersort.stanford.edu/) and run with default parameters and quantile normalization disabled. The LM22 signature was used for inferring cellular proportions. For visualization purposes, different cell subtypes are combined: mast cells, eosinophils, and neutrophils are represented as granulocytes; and monocytes and macrophages are represented as mono/macrophages.

### GeoMx analysis

NanoString GeoMx gene expression analysis using the cancer transcriptome array probe set was performed by the Technology Access Program at NanoString using methods, as previously described *(29*). Briefly, a 5 μm section of FFPE melanoma was dewaxed and stained overnight for DNA, melanocytes (PMEL), epithelia (pan-cytokeratin), and immune cells (CD45) to define areas of interest on the NanoString GeoMx instrument for transcriptional analysis using the human cancer transcriptome array probe set. Twenty-nine ROIs representing five morphological sites (melanoma in situ, invasive tumor, invasive tumor margin, brisk tumor-infiltrating lymphocytes, and exophytic melanoma) were selected. All sample processing and sequencing were performed by the Technology Access Program at NanoString. Tissue images, probe measurements (**Table S4**), and quality control data were provided by NanoString, then analyzed using DESeq2 for differential expression analysis.

### Comparison of Pick-Seq and GeoMx

Similar, but not identical, regions were assayed using GeoMx to compare with Pick-Seq (differences in the areas sampled precluded a one-to-one comparison). Differential gene expression analysis using DESeq2 compared regions of exophytic melanoma and melanoma in situ. Differentially expressed genes with an FDR < 0.05 were compared between Pick-Seq and GeoMx. As the dynamic range of expression scales between the two methods was largely different, we sought to compare them by directional concordance (i.e. up/downregulation) rather than absolute correlation in fold change. Of the 94 common differentially expressed genes, seventeen genes were not directionally concordant. However, only two of those seventeen showed a log-fold change of >1, suggesting that the 15 others might represent low expression noise. Together, this analysis suggested that differential gene expression analysis by Pick- Seq and GeoMx was largely concordant.

## ACKNOWLEDGEMENTS

We thank Jerry Lin for scientific advice and technical assistance and NanoString Inc. for the acquisition of GeoMx DSP data via its Technology Access Program.

## Author contributions

Study design: ZM, YAC, AJN

Quantitative data analysis: AJN, YAC

Data collection: ZM, NGE, LU, RP, JC, YAC, SAB

Pathology analysis and image review: CGL, GFM, SS

Supervision: DMW, EPK, SS, PKS

Writing: ZM, AJN, AAC, PKS

Manuscript was read and approved by all authors.

## Data and materials availability

Raw sequence data and processed counts table can be accessed from NCBI’s GEO repository under accession GSE158564. All code has been deposited on GitHub at https://github.com/sorgerlab.

## SUPPLEMENTARY MATERIALS

**Fig. S1:**
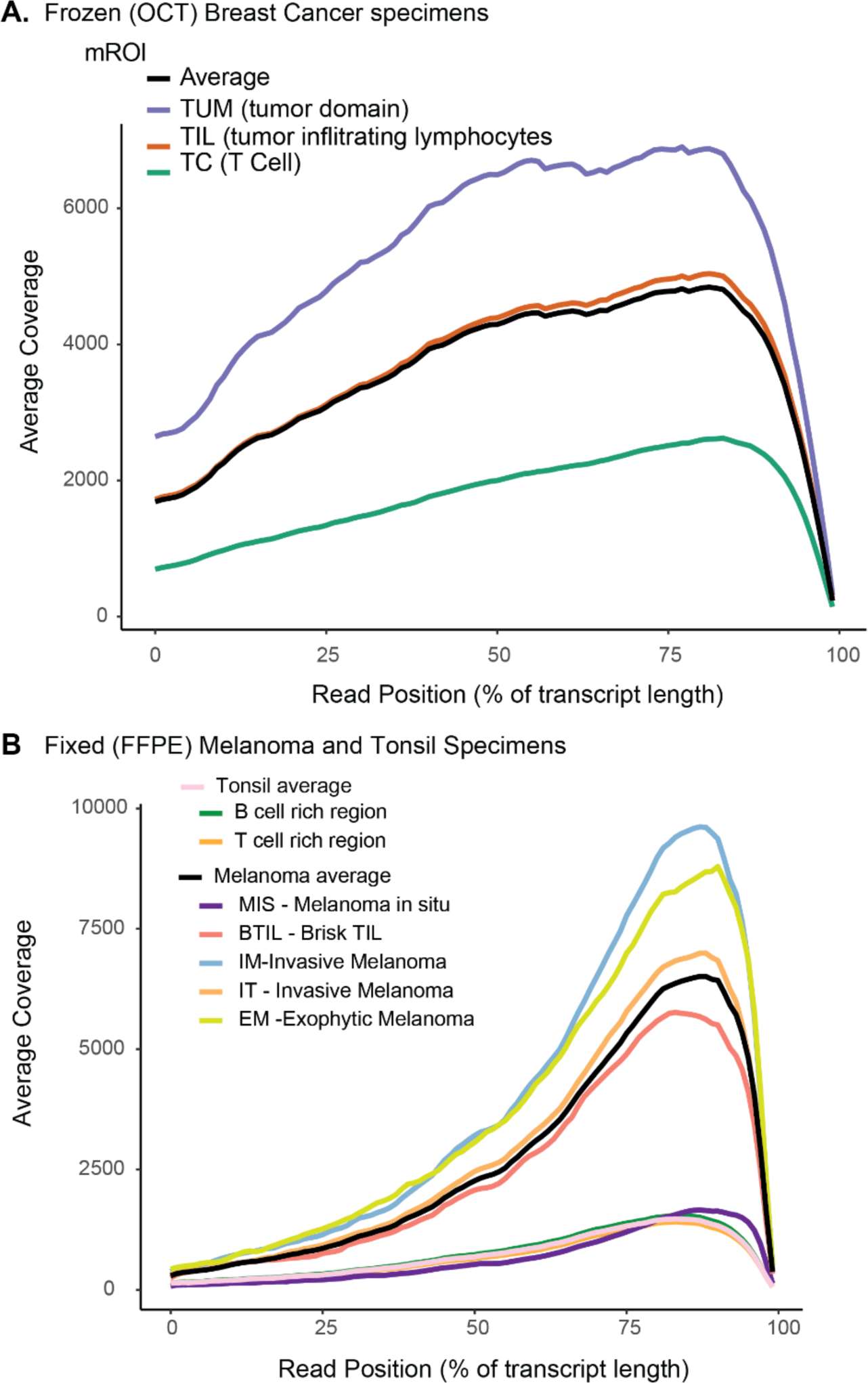
Positions of sequence reads. Plots show mapped-read depth (y-axis) at each relative transcript position (x-axis). The read-depth is averaged across mROI’s belonging to the same sample group. The overall average for each tissue is also shown. (**A)** Frozen (OCT) breast cancer mROIs. (**B)** FFPE tonsil and melanoma mROIs.

**Fig. S2:**
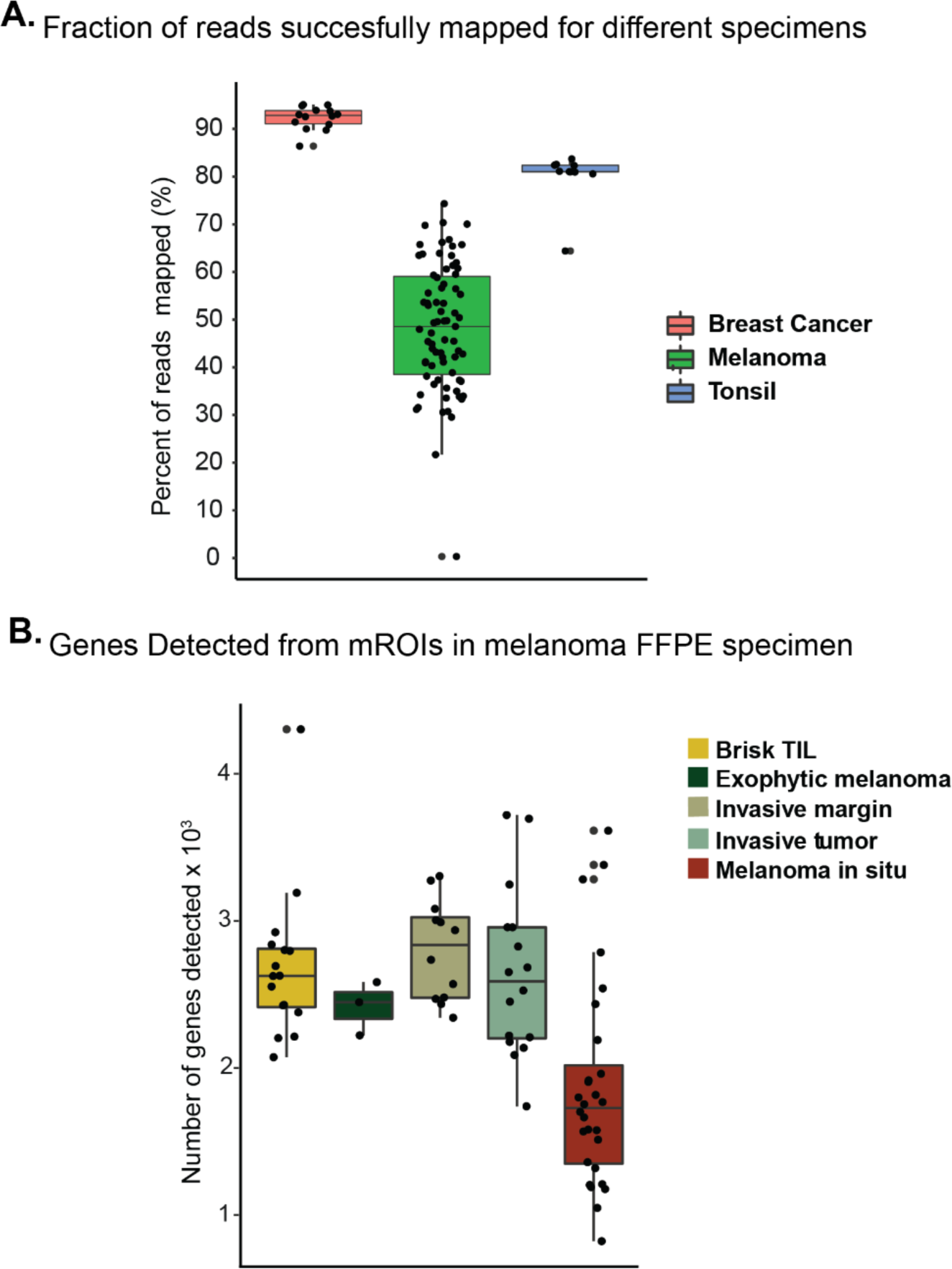
Reads mapped and genes detected for mROIs. **(A**) Boxplot shows the median percentage of reads that could be mapped to the reference genome; the lower and upper hinges correspond to the first and third quartiles. The points represent values for individual mROI’s. (**B)** Number of genes detected in each mROI sorted by histology feature; brisk TIL (n=16), exophytic melanoma (n=3), invasive margin (n=12), invasive tumor (n=16), and MIS (n=28).

**Table S1: Normalized gene expression data for tonsil**

**Table S2: Normalized gene expression data for breast**

**Table S3: Normalized gene expression data for melanoma**

**Table S4: Normalized gene expression data for NanoString GeoMx**

**Table S5: Antibodies for FFPE tissue staining and imaging**

**Table S6: Regions of interest for transcriptional analysis**

